# The most significant differences between male and female rats regarding psychostimulant self-administration behavior are unrelated to biological sex

**DOI:** 10.1101/2024.11.01.621540

**Authors:** Bryce H Showell, Martin O Job

## Abstract

**Background:** The goals of this study were to 1) validate the MISSING (Mapping Intrinsic Sex Similarities as an Integral quality of Normalized Groups) model for psychostimulant self-administration (SA), and 2) utilize it to explain the inconsistencies in the observation of sex differences in psychostimulant SA.

**Methods:** We allowed male and female Long Evans rats (n = 40) to self-administer methamphetamine METH dose 0.1 mg/kg (male n = 9, female n = 18) and saline (male n = 3, female n = 10) via the intravenous route, FR1 schedule, 6 h per day, 5 days per week for 4 weeks. For the MISSING model, we identified behavioral clusters of males and females using normal mixtures clustering analysis of baseline intake, total intake and total intake normalized-to-baseline intake (NBI), followed by unpaired t-tests to compare clusters and Two-way ANOVA to determine if there were any SEX by cluster interactions. For the current model, we grouped our subjects according to biological sex and compared the above variables using unpaired t-tests. For both models, we employed Two-way repeated measures ANOVA and linear regression analysis to analyze SA time course.

**Results:** For saline and METH SA, there were no sex differences when we compared males and females generally, with sex differences evident only when we compared sexes from distinct clusters. The current model could not explain the inconsistencies in the observability of sex differences in METH SA.

**Conclusions:** We validated the MISSING model -it can explain the inconsistencies around sex differences in METH SA.

## Introduction

The epidemic of Methamphetamine (METH) use disorders affects both men and women necessitating efforts to understand sex differences as part of the solution to address this health crisis. These efforts employ sex as a biological variable (SABV) (Clayton, 2016; Miller et al., 2017; Zucker and Beery, 2019) to allow comparisons between males and females (Figure 1A). While there is evidence for sex differences in METH consumption and METH-related behavior, there is also evidence for sex similarities, for review see (Daiwile et al., 2022a). With regards to METH self-administration (SA), most, but not all, studies have observed a lack of sex differences (Table 1). To advance the field, it will be important to be able to explain these inconsistencies (Table 1).

**Figure 1:**
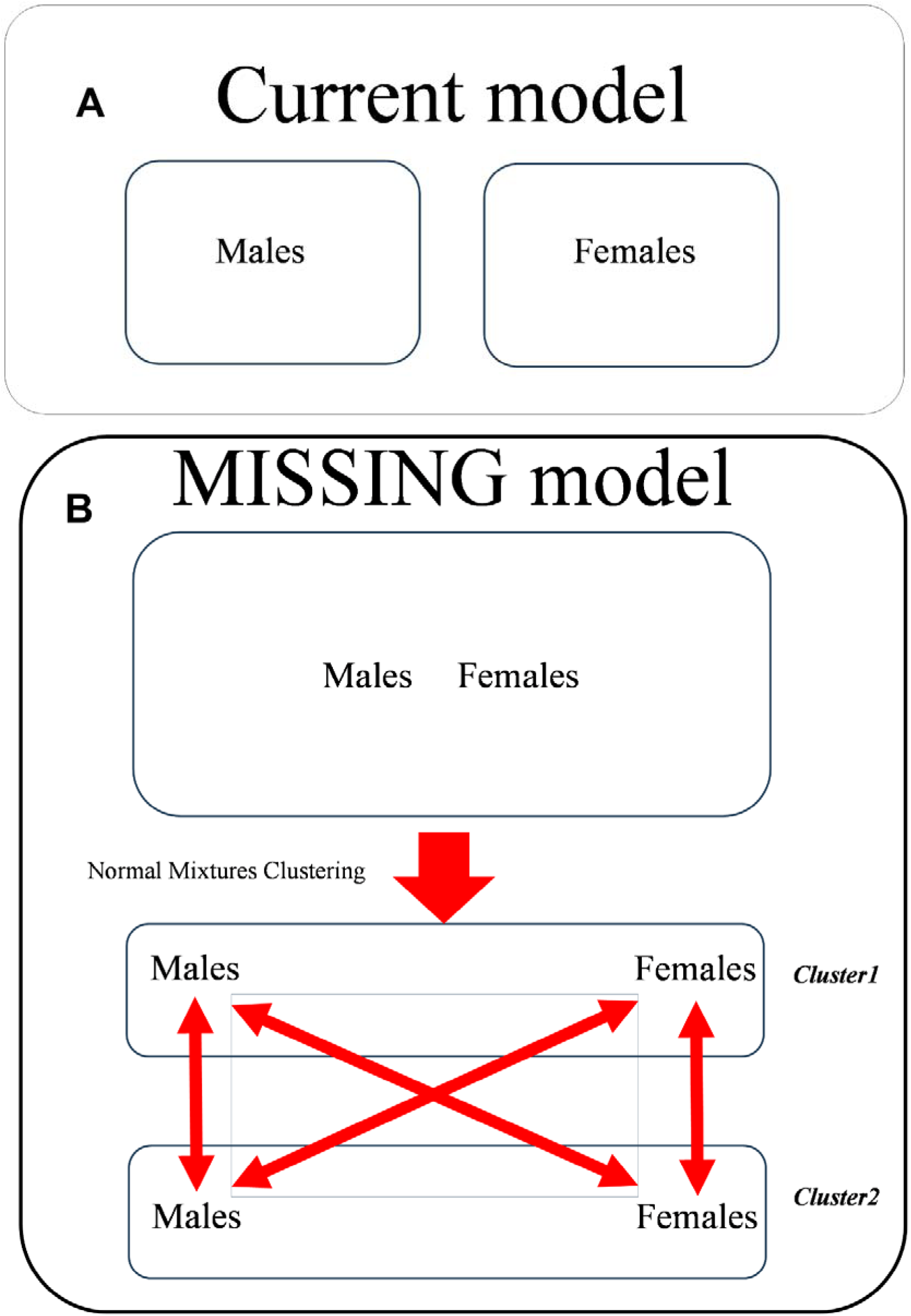
Defining the current and the MISSING models. Fig A represents the current model in which subjects are grouped (and compared) based on biological sex per se. Fig B represents the MISSING (Mapping Intrinsic Sex Similarities as an Integral quality of Normalized Groups) model. Here, the experimenter considers the individual identity (individual similarities/differences from others) of the subject to locate this subject within a behavioral group before the experimenter considers the biological sex of the subject. MISSING involves 1) normal mixtures clustering of all individuals, 2) confirmation that these groups are distinct and then 3) analysis of sex differences within a cluster. The MISSING model proposes that sex differences, are not typically observed in behavior (the vertical arrows within same clusters), except when males and females from different clusters are compared (the diagonal arrows between different sexes in different clusters) – and in this case, the differences observed are not actually biological sex differences but are rather behavioral cluster-related differences.

**Table.**
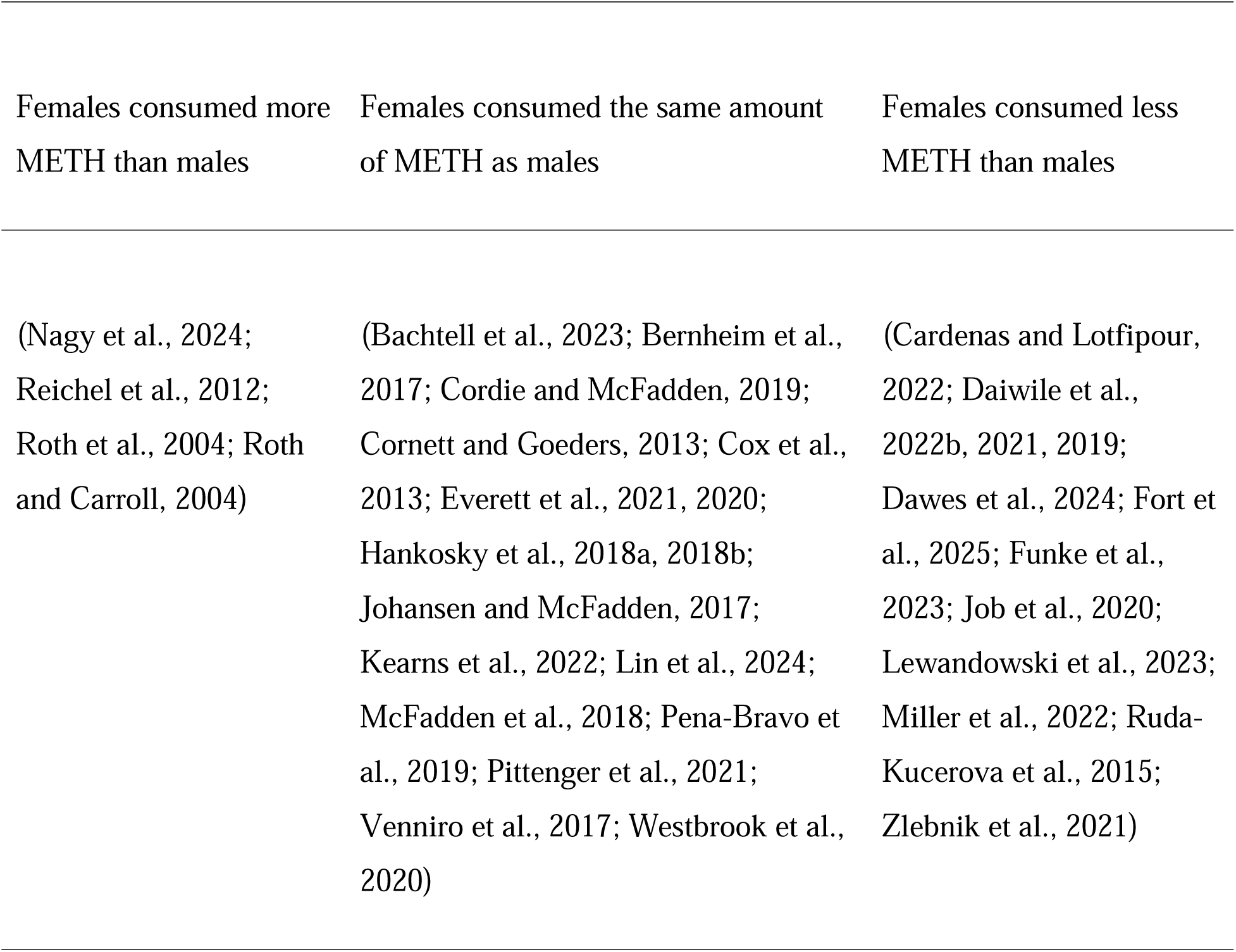
Results from several studies that examined sex differences in methamphetamine self-administration (METH SA) in rats. The columns 1-3 show studies where, relative to males, females consumed more, the same or less METH.

In addition to biological sex differences, males and females are also individuals who can be similar or different from other individuals in the same/different biological sex group. The interplay between biological sex and individual differences/similarities (DS) reveal some interesting observations. For example, some studies of METH SA have designated METH consuming individuals, whether they be males or females, as high versus low METH taker groups (Daiwile et al., 2022b, 2021, 2019; Miller et al., 2022), see also (Everett et al., 2020). While the low takers took less METH than the high takers, as expected, there were no significant differences in METH intake when we compared males versus females within the same behavioral groups (males versus females for low takers, males versus females for high takers) (Daiwile et al., 2022b, 2021, 2019; Miller et al., 2022), see also (Everett et al., 2020). In other words, we can observe sex similarities/differences, depending on what behavioral group(s) of males and females we compare. Thus, the interplay between biological sex and individual DS may be driving the inconsistencies in the observation of sex differences (Table 1), but this is not clear.

The MISSING (Mapping Intrinsic Sex Similarities as an Integral quality of Normalized Groups) model (see Figure 1B) may be able to explain the seemingly disparate findings in Table 1 because it accounts for both the phenomena of individual DS and biological sex. We have validated the MISSING model for psychostimulant-induced locomotor activity in male and female rats following systemic cocaine injection and intra-nucleus accumbens core dopamine injections (Job, 2024; Tigano and Job, 2024). However, we have not validated the MISSING model as a tool for understanding sex differences in the psychostimulant SA paradigm.

There were two goals for this study. The first goal was to validate the MISSING model for psychostimulant SA paradigm, as we previously did for psychostimulant-induced locomotor activity (Job, 2024; Tigano and Job, 2024). The criteria for model validation is that it must confirm, for a psychostimulant SA experiment, the three principles (1-3) of the MISSING model mentioned in the previous reports. The second goal of this study was to utilize the MISSING model as a tool to explain the inconsistencies in the observation of sex differences in METH SA (Table 1).

We conducted METH (and vehicle control) SA experiments in male and female Long Evans rats for 4 weeks. We employed drug SA variables similar to variables we used in previous studies (Job, 2024; Tigano and Job, 2024). For the MISSING model, we conducted normal mixtures clustering analysis of these variables for every individual, regardless of biological sex, to identify behavioral clusters prior to conducting comparisons between males and females (Figure 1B). For the current model (SABV) we compared these variables using biological sex to group our subjects (Figure 1A). Our methods, results and discussions are below.

## Methods and Materials

### Animals

All procedures and treatments were approved by the National Institute on Drug Abuse Animal Care and Use Committee and followed the guidelines outlined in the National Institutes of Health (NIH) *Guide for the Care and Use of Laboratory Animals*. The animals, male and female Long Evans, were obtained from the National Institute on Drug Abuse (NIDA).

### Experiments

Rats were single housed on a 12-hour reversed light/dark cycle with free access to food and water. For this current analysis, a total of forty (n = 40) subjects were employed. The breakdown of animal utilization was as follows: eight male (n = 9) and sixteen female (n = 18) were used for METH SA studies and three male (n = 3), and ten females (n = 10) were used as saline controls. The rats were adults, and the female and male rats weighed within the range of 250-450 and 500–650 grams, respectively, on the first day of the SA experiments. For jugular vein catheterization surgery, see (Job et al., 2020). All animals were allowed to recover from surgery for approximately one week before SA procedures were initiated. After rats had recovered from the surgical procedures described above, the rats were allowed to SA METH (0.1 mg/kg/infusion, or saline in control rats) on the FR1 schedule for 6 h per session for 5 days a week (no SA experiments on the weekend) for 4 weeks, as described in (Job et al., 2020).

### Variables

We employed variables similar to the ones we used to validate the MISSING model for psychostimulant-induced locomotor activity, see the variables: baseline activity, drug activity and drug activity normalized-to-baseline activity (NBA) in the previous reports (Job, 2024; Tigano and Job, 2024). To adapt these variables for SA studies, we estimated baseline intake, total intake and total intake normalized-to-baseline intake (NBI). These variables are shown in Figure 2 and defined below. Baseline intake was defined as the total amount of drug consumed (number of active lever presses/ number of infusions) by the end of the first week of a 4-week drug SA session. Total intake was defined as the total drug consumed throughout the 4-week drug SA experiment. Total intake NBI was calculated as below:

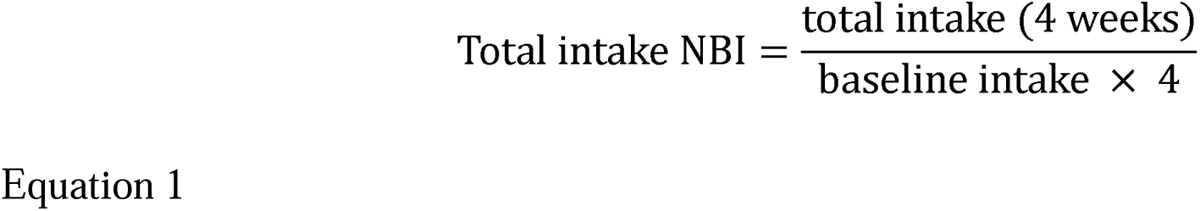

**Figure 2:**
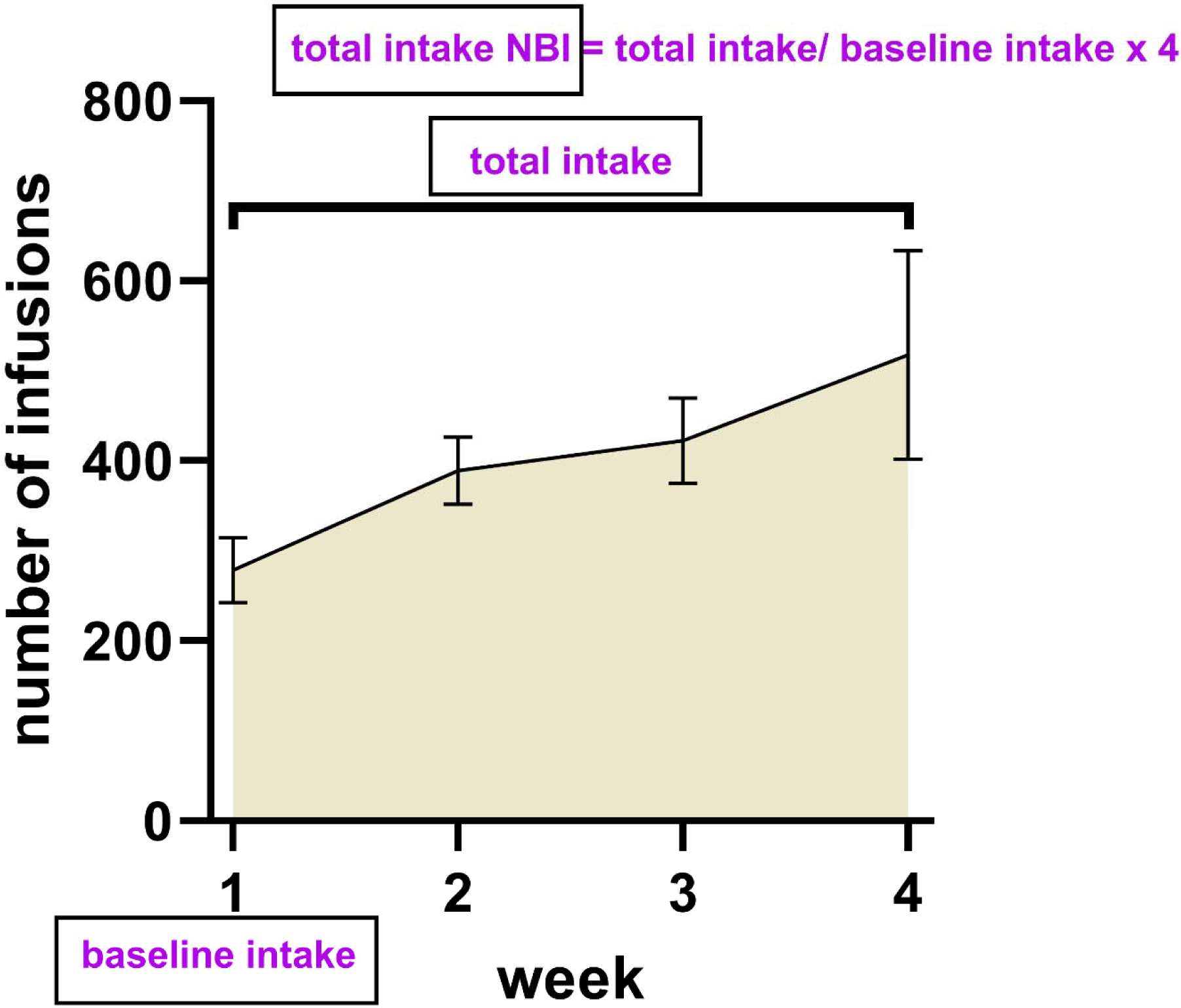
Variables. We employed three variables which we used in cluster analysis to identify our groups: baseline intake, total intake and total intake normalized-to-baseline intake (total intake NBI). These variables are similar to the variables we used in previous studies (baseline activity, drug activity, drug activity normalized-to-baseline activity) to validate the MISSING model for psychostimulant-induced locomotor activity (Job, 2024; Tigano and Job, 2024). But because this study is for psychostimulant self-administration, we had to adapt the variables we utilized by adjusting them slightly. Baseline intake is defined as the number of infusions obtained in the first week of the four-week SA sessions. Total intake is defined as the total number of infusions over the 4 weeks (sum of all drug infusions for the duration of the experiments). Total intake NBI was calculated as a function of the baseline intake and is expressed mathematically as total intake divided by (baseline intake x 4) see equation 1 in the Methods section. Total intake NBI expresses a fold change of drug intake over baseline intake (this is a normalized variable).

### Statistical analysis

GraphPad Prism v 10 (GraphPad Software, San Diego, CA), SigmaPlot 14.5 (Systat Software Inc., San Jose, CA) and JMP Pro v 17 (SAS Institute Inc., Cary, NC) were employed for statistical analysis. Data were expressed as mean ± SEM. For the current model, we grouped our subjects using biological sex only and, using unpaired t-tests, we compared the following variables (baseline intake, total intake, total intake NBI) for saline and METH SA groups. For the MISSING model, we conducted normal mixtures clustering analysis of the variables mentioned above to identify distinct clusters, if any. We employed linear regression analysis and unpaired t-tests (or One-way ANOVA) to establish that these clusters were different. Identification of clusters was followed by a Two-way ANOVA to determine if there was a SEX (males, females) by cluster interaction and main effects of SEX and of cluster. With biological sex groups and behavioral clusters identified, we conducted analysis of the time course of drug consumption using a Two-way repeated measures ANOVA to determine if there were SEX × time or cluster × time interactions, respectively. We also employed linear regression for time course data analysis. Statistical significance was set at P < 0.05 for all analyses with Tukey’s post hoc test employed when significance was detected.

## Results

### Identification of clusters for the METH SA group (MISSING model)

For the METH SA group, normal mixtures clustering analysis of baseline intake, total intake and total intake NBI for all subjects (n = 27), including males and females, yielded two clusters which we termed cluster1 and cluster2. Cluster1 (n = 15) consisted of n = 5 males and n = 10 females whereas cluster2 (n = 12) consisted of n = 4 males and n = 8 females (Figure 3A-B).

**Figure 3:**
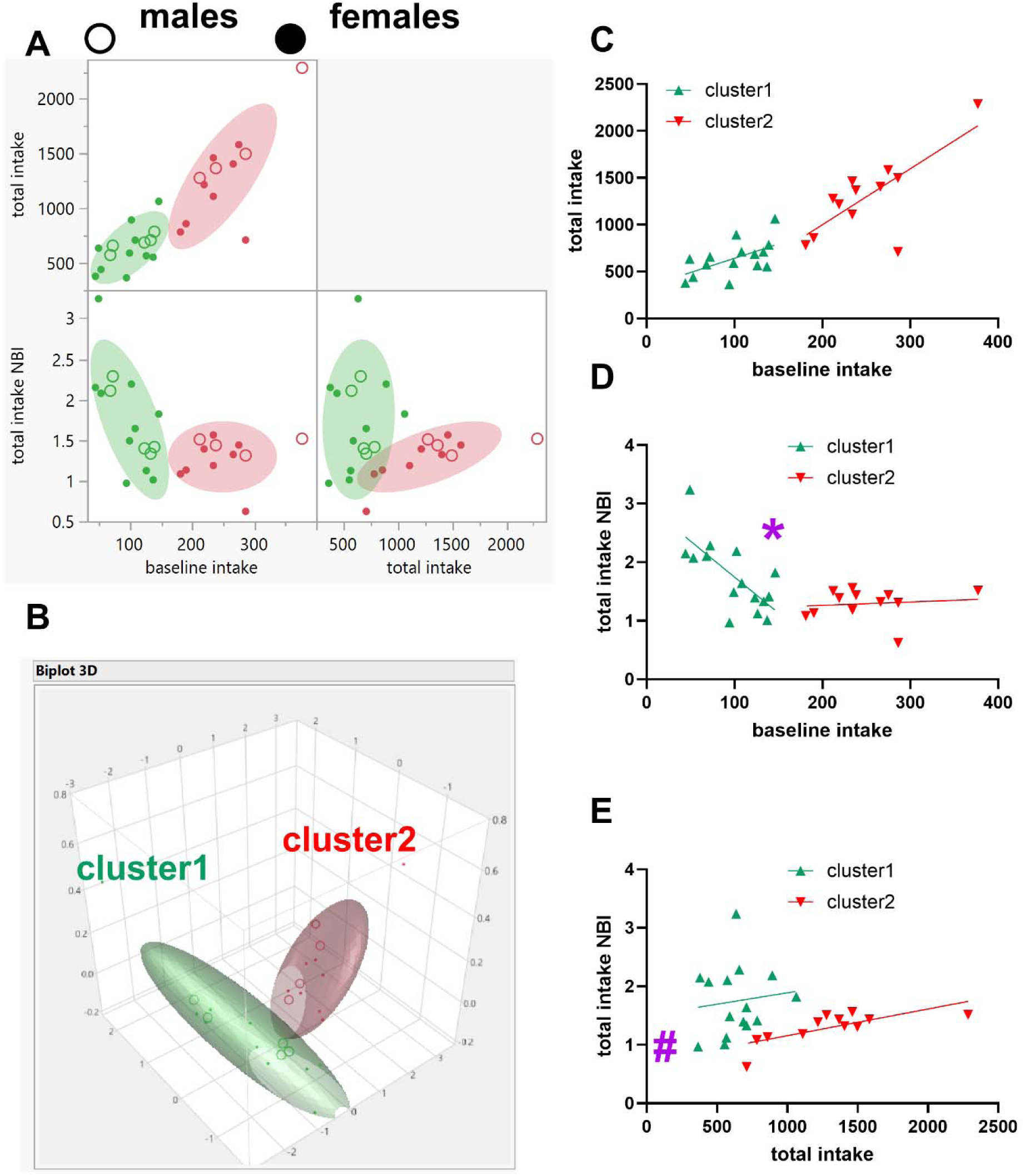
Identification of clusters for the METH SA group (MISSING model). The open circles represent males while the closed circles represent females. Normal mixtures clustering analysis of baseline intake, total intake and total intake NBI for all subjects (n = 27), including males and females, yielded two clusters which we termed cluster1 and cluster2. Cluster1 (n = 15) consisted of n = 5 males and n = 10 females whereas cluster2 (n = 12) consisted of n = 4 males and n = 8 females (Fig A-B). The relationships between baseline intake and total intake were similar for both clusters (Fig C). The relationship between baseline intake and total intake NBI were different for these clusters for slope (Fig D) and the relationship between total intake and total intake NBI were different for these clusters for y-axis intercept (Fig E). In summary, normal mixtures clustering identified two distinct clusters each consisting of males and females.

The relationships between baseline intake and total intake (Figure 3C) were significant for both cluster1 (F 1, 13 = 6.369, P = 0.0254, R^2^ = 0.33) and cluster2 (F 1, 10 = 12.51, P = 0.0054, R^2^ = 0.56). Comparisons of these relationships showed no differences between cluster1 and cluster2 with regards to slope (F 1, 23 = 1.807, P = 0.1920) and y-axis intercept (F 1, 24 = 0.2087, P = 0.6519). The relationships between baseline intake and total intake NBI (Figure 3D) were significant for cluster1 (F 1, 13 = 13.13, P = 0.0031, R^2^ = 0.50) but not for cluster2 (F 1, 10 = 0.1361, P = 0.7199, R^2^ = 0.013), and comparisons of these relationships showed significant differences between cluster1 and cluster2 with regards to slope (F 1, 23 = 12.69, P = 0.0017) and y-axis intercept (P = 0.0021). The relationships between total intake and total intake NBI (Figure 3E) were significant for cluster2 (F 1, 10 = 12.09, P = 0.0059, R^2^ = 0.55) but not for cluster1 (F 1, 13 = 0.1871, P = 0.6724, R^2^ = 0.014), and comparisons of these relationships showed no differences between cluster1 and cluster2 with regards to slope (F 1, 23 = 0.007335, P = 0.9325) but significant difference with regards to the y-axis intercept (F 1, 24 = 7.660, P = 0.0107).

### Model comparisons: average of variables (METH SA group)

The baseline intake, total intake and total intake NBI for males in the METH SA group (n = 9) were 183.11 ± 34.50 infusions, 1093.56 ± 188.66 infusions and 1.589 ± 0.118, respectively. The baseline intake, total intake and total intake NBI for METH SA group females (n = 18) were 157.94 ± 18.90 infusions, 851.06 ± 88.97 infusions and 1.524 ± 0.143, respectively. There were no significant differences (unpaired t-tests) between males and females with regards to baseline intake (P = 0.4916, Figure 4A), total intake (P = 0.1955, Figure 4B) and total intake NBI (P = 0.7695, Figure 4C).

**Figure 4:**
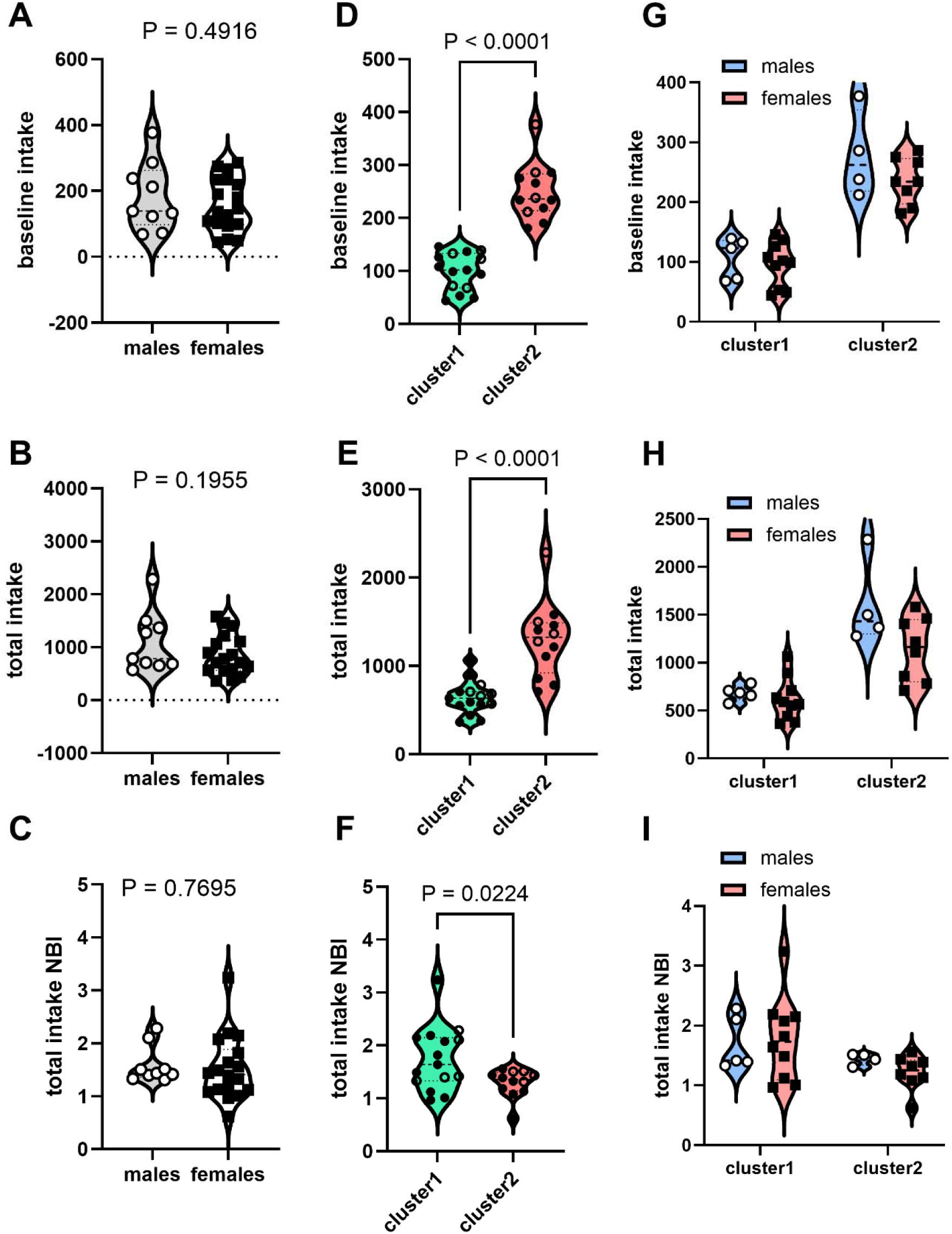
Current versus MISSING model comparisons: average of variables (METH SA group). Unpaired t-tests determined that there were no differences between males and females with regards to baseline intake (Fig A), Total intake (Fig B) and Total intake NBI (Fig C). Unpaired t-tests determined that there were differences between cluster1 and cluster2 with regards to baseline intake (Fig D), Total intake (Fig E) and Total intake NBI (Fig F). Two-way ANOVA with factors being SEX (males, females) and cluster (cluster1, cluster2) revealed no SEX by cluster interaction for any of the variables (Fig G-I). For baseline intake, there was a main effect of cluster but not SEX. For total intake, the analysis revealed a main effect of SEX and cluster, with the effects of cluster (P < 0.0001) being more significant than the effects of SEX (P = 0.0337). For total intake NBI, the analysis revealed no main effects of SEX and cluster (P > 0.05). In summary, the current model did not detect any significant differences between males and females while the MISSING model detected significant differences between clusters of males and females. The differences between clusters exceeded the differences between biological sex – females in one cluster were similar to males in the same cluster but were different from males and females in a different cluster.

The mean ± SEM for baseline intake, total intake and total intake NBI, respectively, for cluster1 were as follows: 99.53 ± 9.04, 640.27 ± 47.87 and 1.749 ± 0.156. For cluster2, baseline intake, total intake and total intake NBI, respectively, were 249.83 ± 15.33, 1296. 42 ± 122.67 and 1.291 ± 0.076. There were significant differences between cluster1 and cluster2 with regards to these same variables: baseline intake (P < 0.0001, Figure 4D), total intake (P < 0.0001, Figure 4E) and total intake NBI (P = 0.0224, Figure 4F).

Because these clusters consist of both males and females, we wanted to understand if there were any SEX (males, females) × cluster (cluster1, cluster2) interactions. For baseline intake (Figure 4G), Two-way ANOVA did not reveal a SEX × cluster interaction (F 1, 23 = 0.7841, P = 0.3851) or a main effect of SEX (F 1, 23 = 2.300, P = 0.1430) but did reveal a main effect of cluster (F 1, 23 = 76.83, P < 0.0001). Likewise, for total intake (Figure 4H), Two-way ANOVA did not reveal a SEX × cluster interaction (F 1, 23 = 2.949, P = 0.0994) but did reveal main effects of SEX (F 1, 23 = 5.101, P = 0.0337) and cluster (F 1, 23 = 38.04, P < 0.0001). For total intake NBI (Figure 4I), Two-way ANOVA did not reveal a SEX × cluster interaction (F 1, 23 = 0.4975, P = 0.4877) or a main effect of SEX (F 1, 23 = 0.1568, P = 0.6958) or cluster (F 1, 23 = 3.979, P = 0.0580), but note that the main effect of cluster was almost significant.

### Model comparisons: average of variables (saline SA group)

The baseline intake, total intake and total intake NBI for the saline SA group males (n = 3) were 45.00 ± 0.58 infusions, 184.33 ± 6.96 infusions and 1.025 ± 0.045, respectively. The baseline intake, total intake and total intake NBI for females in the saline SA group (n = 10) were 69.30 ± 6.63 infusions, 212.80 ± 10.85 infusions and 0.802 ± 0.044, respectively.

For the saline SA group, there were no significant differences (unpaired t-tests) between males and females with regards to baseline intake (P = 0.0775, Figure 5A), total intake (P = 0.1966, Figure 5B), but there were significant differences with respect to total intake NBI (P = 0.0248, Figure 5C).

**Figure 5:**
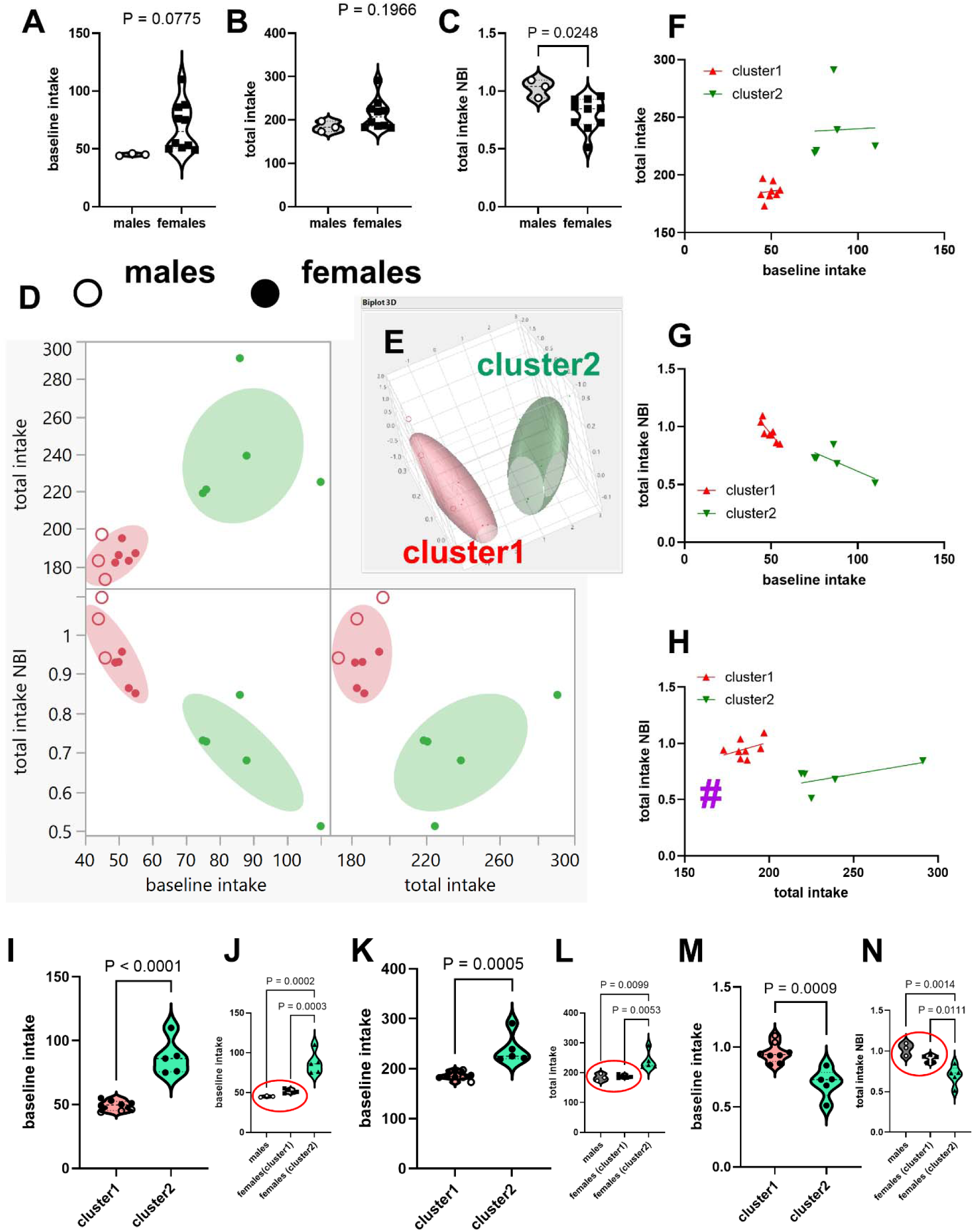
Identification of clusters for the METH SA group (MISSING model) and model comparisons. For saline SA, unpaired t-tests determined that there were no differences between males and females with regards to baseline intake (Fig A), and Total intake (Fig B) but there were significant differences between sexes for Total intake NBI (Fig C). Normal mixtures clustering analysis of baseline intake, total intake and total intake NBI for all subjects (n = 13), including males and females, yielded two clusters which we termed cluster1 and cluster2. Cluster1 (n = 8) consisted of n = 3 males and n = 5 females whereas cluster2 (n = 5) consisted of n = 0 males and n = 5 females (Fig D-E). The relationships between baseline intake and 1) total intake, and 2) total intake NBI were similar for both clusters (Fig F-G). The relationship between total intake and total intake NBI were different for these clusters for y-axis intercept (Fig H). Unpaired t-tests determined that there were differences between cluster1 and cluster2 with regards to baseline intake (Fig I), Total intake (Fig K) and Total intake NBI (Fig M). Because there was a cluster that included only females, we could not conduct a Two-way ANOVA. A One-way ANOVA revealed that the females in cluster2 were different from both the males and females in cluster1 while the males and females in cluster1 were not different for any variable (Fig J, L and N). In summary, the current model did not detect any significant differences between males and females for baseline intake and total intake (but see total intake NBI) while the MISSING model detected significant differences between clusters of males and females for all these variables. The differences between clusters exceeded the differences between biological sex – the differences between females in cluster2 and females in cluster1 exceeded the difference between males and females in cluster1.

Normal mixtures clustering analysis of baseline intake, total intake and total intake NBI for all subjects (n = 13), including males and females, yielded two clusters which we termed cluster1 (n = 3 males, n = 5 females) and cluster2 (n = 0 males, n = 5 females), see Figure 5D-E.

The relationships between baseline intake and total intake (Figure 5F) were not significant for both cluster1 (F 1, 6 = 0.069, P = 0.8011, R^2^ = 0.011) and cluster2 (F 1, 3 = 0.0043, P = 0.9520, R^2^ = 0.0014) and these lines showed no differences with regards to slope (F 1, 9 = 0.0034, P = 0.9545) and y-axis intercept (F 1, 10 = 3.201, P = 0.1039). The relationships between baseline intake and total intake NBI (Figure 5G) were significant for cluster1 (F 1, 6 = 18.21, P = 0.0053, R^2^ = 0.75) but not for cluster2 (F 1, 3 = 3.945, P = 0.1412, R^2^ = 0.57), and comparisons of these relationships showed no significant differences with regards to slope (F 1, 9 = 3.166, P = 0.1089) and y-axis intercept (F 1, 10 = 0.2281, P = 0.6432). The relationships between total intake and total intake NBI (Figure 5H) were not significant for cluster1 (F 1, 6 = 1.130, P = 0.3286, R^2^ = 0.16) and cluster2 (F 1, 3 = 1.930, P = 0.2590, R^2^ = 0.39) and comparisons of these relationships showed no differences with regards to slope (F 1, 9 = 0.1377, P = 0.7191) but significant differences with regards to the y-axis intercept (F 1, 10 = 19.76, P = 0.0012).

Unlike what we obtained for comparisons between males and females, there were significant differences between cluster1 and cluster2 with regards to baseline intake (P < 0.0001, Figure 5I), total intake (P = 0.0005, Figure 5K) and total intake NBI (P = 0.0009, Figure 5M). For the saline SA group, one of the clusters contained males and females while the other cluster consisted of only females (Figure 5D-E) and we could not conduct an analysis to determine if there were SEX (males, females) × cluster (cluster1, cluster2) interactions for all the variables we obtained.

Therefore, we conducted One-way ANOVA which determined significant differences between these three groups for baseline intake (F 2, 10 = 27.37, P < 0.0001, Figure 5J), total intake (F 2, 10 = 10.86, P = 0.0031, Figure 5L) and total intake NBI (F 2, 10 = 13.89, P = 0.0013, Figure 5N). The details were as follows: 1) there were no significant differences between males and females within cluster1 for any of the variables, 2) there were significant differences between males in cluster1 and females in cluster2 for all variables, and 3) there were significant differences between females in cluster1 and females in cluster2 for all variables.

### Model comparisons: saline and METH SA time course

For the saline SA group, we had n = 3 males and n = 10 females. We analyzed the saline intake time course data (Figure 6A) using Two-way repeated measures ANOVA with factors SEX (males, females) and time (weeks 1-4) as independent variables and intake (number of infusions) as the dependent variable. We detected no SEX × time interaction (F 3, 33 = 2.811, P = 0.0545), no main effect of SEX (F 1, 11 = 1.890, P = 0.1966), and no main effect of time (F 1.242, 13.66 = 2.114, P = 0.1674). Because there was no interaction, we did not proceed with post hoc tests.

**Figure 6.**
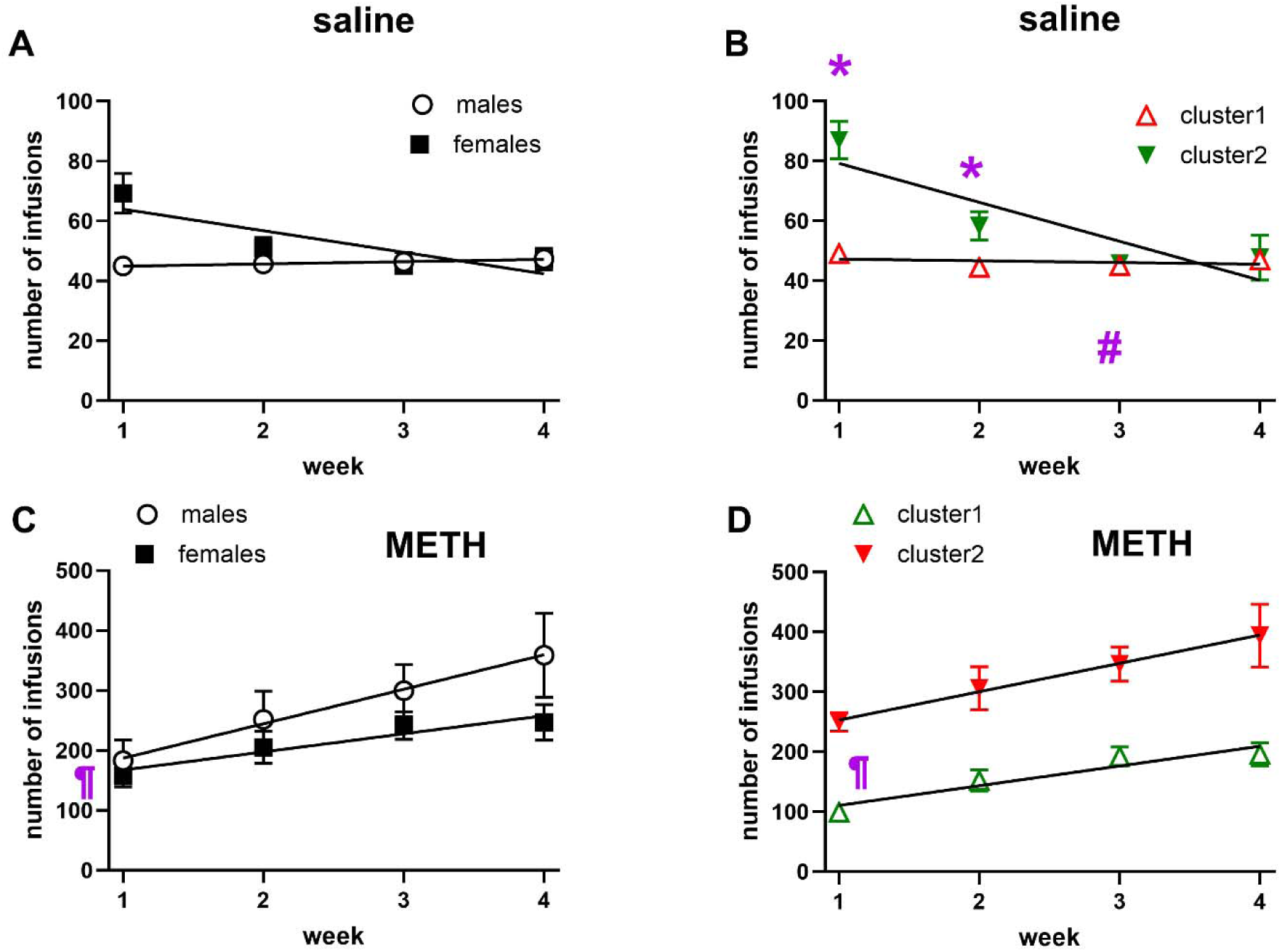
Differences between cluster1 and cluster2 exceeds differences between males and females for saline and METH SA time course. For the saline SA group, the current model had n = 3 males and n = 10 females and for the MISSING model, we had cluster1 (n = 8) and cluster2 (n = 5). Using Two-way repeated measures ANOVA with factors SEX (2 levels: males, females) and time (4 levels: weeks 1-4) and dependent variable being intake (number of infusions), we analyzed the saline intake time course data (Fig A). We detected no SEX × time interaction (P > 0.05), no main effect of SEX (P > 0.05), and no main effect of time (P > 0.05). Interestingly, when we did the same for saline SA clusters (Fig B), we detected a significant cluster × time interaction (P < 0.0001), a significant main effect of time (P = 0.0001), and a significant main effect of cluster (P = 0.0005). For the METH SA group, the current model had n = 9 males and n = 18 females, and the MISSING model had cluster1 (n = 5 males and n = 10 females) and cluster2 (n = 4 males and n = 8 females). We did not detect any SEX × time interaction (P > 0.05), or any main effect of SEX (0.1955), but we detected a main effect of time (P < 0.0001). Interestingly, when we did the same for the clusters, we detected no significant cluster × time interaction (P = 0.4172), but significant main effects of time (P < 0.0001), and cluster (P < 0.0001). Linear regression analysis revealed differences in slope for the saline SA groups and in y-axis intercepts for the METH SA group, but both cases the differences between these variables for cluster1 and cluster2 exceeded the differences between males and females. The * represent differences between clusters and the # represent differences compared to week1 (Tukey’s post hoc tests). The ¶ represents differences between y-axis intercepts (linear regression analysis). In summary, for both saline and METH SA, differences between clusters exceeded differences between biological sex.

We analyzed the time course data of the two saline SA group clusters identified by the MISSING model (Figure 6B): cluster1 had n = 8 subjects (n = 3 males and n = 5 females) and cluster2 had n = 5 subjects (n = 0 males and n = 5 females). Our analysis revealed a significant cluster × time interaction (F 3, 33 = 16.22, P < 0.0001), a significant main effect of time (F 1.294, 14.23 = 22.39, P = 0.0001), and a significant main effect of cluster (F 1, 11 = 23.82, P = 0.0005). Tukey’s post hoc tests revealed differences between clusters on week1 (P = 0.0031) and week2 (P = 0.04180 (Figure 6B) and between week1 and week3 for cluster2 (P = 0.0198).

For the METH SA group, we had n = 9 males and n = 18 females. We did not detect any SEX × time interaction (F 3, 75 = 2.424, P = 0.0724), no main effect of SEX (F 1, 25 = 1.769, P = 0.1955), but a main effect of time (F 2.283, 57.06 = 23.04, P < 0.0001). Because there was no interaction (Figure 6C), we did not proceed with post hoc tests.

For the METH SA group, the MISSING model identified two clusters: cluster1 had n = 15 subjects (n = 5 males and n = 10 females) and cluster2 had n = 12 subjects (n = 4 males and n = 8 females). Analysis of the time course data using Two-way repeated measures ANOVA (Figure 6D) revealed no significant cluster × time interaction (F 3, 75 = 0.9579, P = 0.4172), but significant main effects of time (F 2.212, 55.30 = 20.44, P < 0.0001), and cluster (F 1, 25 = 29.08, P < 0.0001). Because there was no interaction (Figure 6D), we did not proceed with post hoc tests.

We also conducted linear regression analysis of the time course data to estimate the slopes and y-axis intercepts and determine differences (between groups) for these variables, if any. There were differences in slope for the saline SA groups for both males versus females (F 1, 48 = 4.654, P = 0.0360, Figure 6A) and cluster1 versus cluster2 (F 1, 48 = 27.31, P < 0.0001, Figure 6B). For the METH SA group, there were no differences with regards to the slope for males versus females (F 1, 104 = 1.540, P = 0.2173, Figure 6C) and for cluster1 versus cluster2 (F 1, 104 = 0.8011, P = 0.3728 Figure 6D). There were differences in the y-axis intercept(s) for the METH SA groups for both males versus females (F 1, 105 = 6.044, P = 0.0156, Figure 6C) and cluster1 versus cluster2 (F 1, 105 = 83.42, P < 0.0001, Figure 6D).

### Comparisons of males and females within and between clusters (time course data)

Because these METH SA clusters consisted of both males and females, we wanted to compare the time course data for METH consumption for males and females within the same cluster (Figure 7A-B) and between different clusters (Figure 7C-D). For this analysis, we utilized Two-way repeated measures ANOVA with SEX (males, females) and time (4 weeks) as factors. All comparisons revealed the main effect of time (P < 0.05).

**Figure 7:**
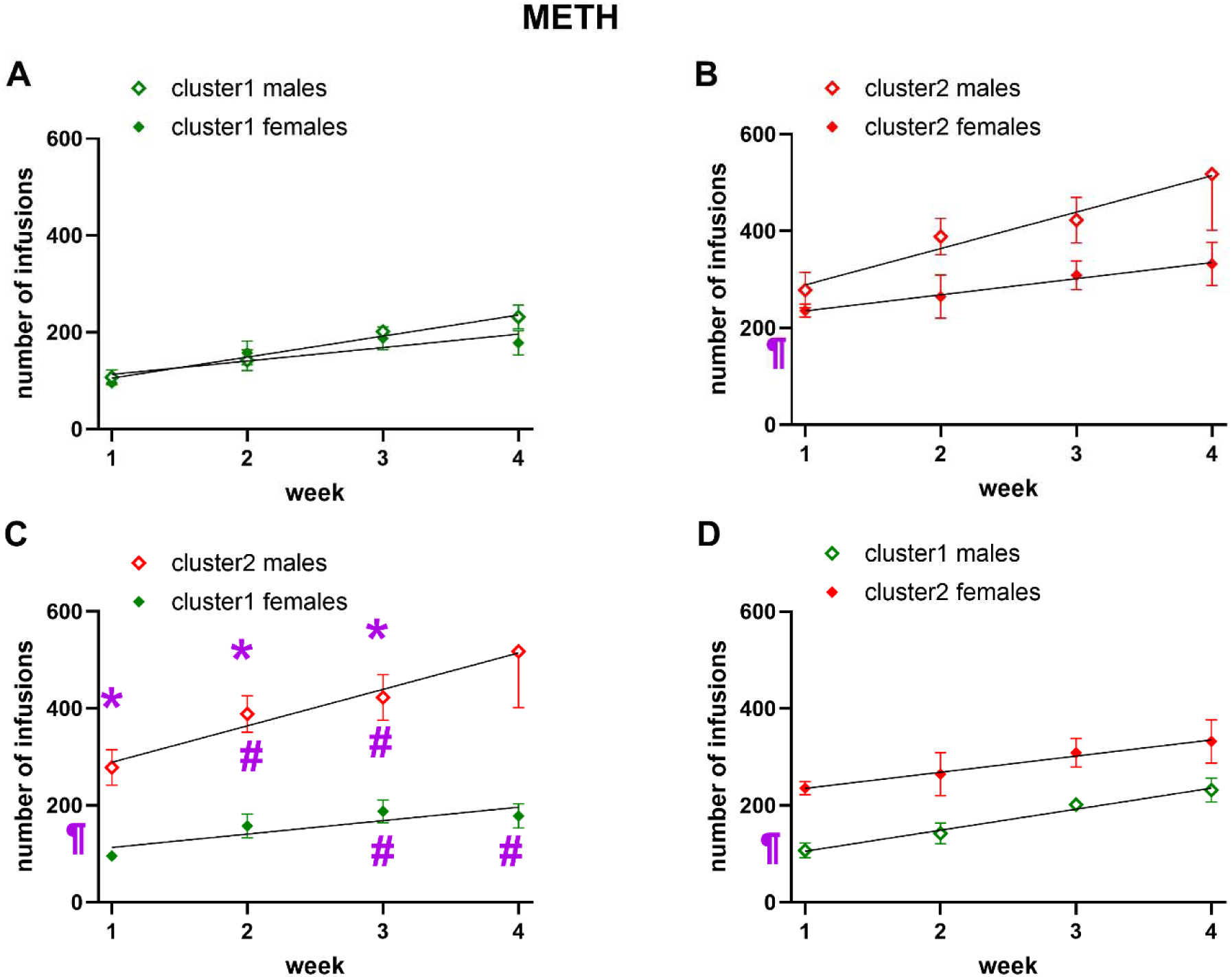
Differences between males and females regarding methamphetamine self-administration are not due primarily to biological sex. Table 1 shows that there are inconsistencies in the observability of sex differences in METH SA with females consuming more, same or less METH than males. The MISSING model identified 2 clusters consisting of males and females: cluster1 (n = 5 males and n = 10 females) and cluster2 (n = 4 males and n = 8 females). We analyzed the time course data males versus females for the different clusters using Two-way repeated measures ANOVA with factors being SEX (males, females) and time (weeks 1-4). All comparisons revealed the main effect of time (P < 0.05). For cluster1 males versus cluster1 females (Fig A), there was no SEX × time interaction (P = 0.3035), and no main effect of SEX (P = 0.5516). Likewise, for cluster2 males versus cluster2 females (Fig B), there was no SEX × time interaction (P = 0.0.1635), and no main effect of SEX (P = 0.0698). For cluster2 males versus cluster1 females (Fig C), there was a significant SEX × time interaction (P = 0.0220), and a main effect of SEX (P = 0.0001). For cluster1 males versus cluster2 females (Fig D), there was no significant SEX × time interaction (P = 0.9227), but there were main effects of SEX (P = 0.0122). We also conducted linear regression analysis of the time course data to estimate the slopes and y-axis intercepts and determine sex differences in these variables, if any. There were differences in slope only for cluster2 males versus cluster1 females (Fig C). The * represent differences between SEX and the # represent differences compared to week1 (Tukey’s post hoc tests, Two-way repeated measures ANOVA). The ¶ represents differences between y-axis intercepts (linear regression analysis). Note that there were no differences in y-axis intercept between males and females from the same cluster (cluster1, Fig A). Even when there were differences in y-axis intercept between males and females in the same cluster (cluster2, Fig B), these differences were not as significant as the differences in y-axis intercept between males and females from different clusters (Fig C-D), confirming the MISSING model principles. Our results can explain Table 1 by showing that, depending on the clusters compared, females can consume more (Fig D), same (Fig A) and less (Fig B-C) than males.

For cluster1 males versus cluster1 females (Figure 7A), Two-way repeated measures ANOVA revealed that there was no SEX × time interaction (F 3, 39 = 1.254, P = 0.3035) and no main effect of SEX (F 1, 13 = 0.3736, P = 0.5516).

Likewise, for cluster2 males versus cluster2 females (Figure 7B), there was no SEX × time interaction (F 3, 30 = 1.827, P = 0.0.1635) and no main effect of SEX (F 1, 10 = 4.121, P = 0.0698).

For cluster2 males versus cluster1 females (Figure 7C), there was a significant SEX × time interaction (F 3, 36 = 3.623, P = 0.0220) and a main effect of SEX (F 1, 12 = 30.93, P = 0.0001). Tukey’s post hoc tests revealed differences between clusters on week1 (P = 0.0111), week2 (P = 0.0023), and week3 (P = 0.0082) with week4 almost significantly different (P = 0.0579). Tukey’s post hoc tests also revealed differences for week1 versus 1) week2 (P = 0.0077) and 2) week3 (P = 0.0067) for cluster2 males while we detected differences for cluster1 females when we compared week1 versus 1) week3 (P = 0.0076) and 2) week4 (P = 0.0477).

For cluster1 males versus cluster2 females (Figure 7D), there was no significant SEX × time interaction (F 3, 33 = 0.1597, P = 0.9227), but there was a main effect of SEX (F 1, 11 = 8.980, P = 0.0122).

We also conducted linear regression analysis of the time course data to estimate the slopes and y-axis intercepts and determine differences between groups for these variables, if any. For cluster1 males versus cluster1 females (Figure 7A), there were no differences in the slope (F 1, 56 = 1.070, P = 0.3054) or y-axis intercept (F 1, 57 = 0.8774, P = 0.3529). For cluster2 males versus cluster2 females (Figure 7B), there were no differences in the slope (F 1, 44 = 2.029, P = 0.1614), but differences in y-axis intercept (F 1, 45 = 12.31, P = 0.0010). For cluster2 males versus cluster1 females (Figure 7C), there were significant differences in the slope (F 1, 52 = 4.075, P = 0.0487). For cluster1 males versus cluster2 females (Figure 7D), there were no differences in the slope (F 1, 48 = 0.2393, P = 0.6270), but differences in y-axis intercept (F 1, 49 = 25.51, P < 0.0001).

## Discussion

One of the goals of this study was to validate the MISSING model for psychostimulant SA as we have previously done for psychostimulant-induced locomotor activity (Job, 2024; Tigano and Job, 2024). We show that 1) there are no sex differences when we compare males and females within the same behavioral group for METH/saline SA, 2) sex differences occur when we compare males and females from different groups, and 3) even if/when we detect sex differences between males and females in the same behavioral group, these differences will not exceed the differences between males and females from different behavioral groups. Our results validated the MISSING model for METH SA.

The other goal was to utilize the MISSING model as a tool to explain the inconsistencies in the observability of sex differences in METH SA. By showing that there are no sex differences within clusters but there are sex differences when males and females from different clusters are compared, our data can explain all the different observations including where 1) females > males (Nagy et al., 2024; Reichel et al., 2012; Roth et al., 2004; Roth and Carroll, 2004), 2) females < males (Cardenas and Lotfipour, 2022; Daiwile et al., 2022b, 2021, 2019; Dawes et al., 2024; Fort et al., 2025; Funke et al., 2023; Job et al., 2020; Lewandowski et al., 2023; Miller et al., 2022; Ruda-Kucerova et al., 2015; Zlebnik et al., 2021), and 3) females = males (Bachtell et al., 2023; Bernheim et al., 2017; Cordie and McFadden, 2019; Cornett and Goeders, 2013; Cox et al., 2013; Everett et al., 2021, 2020; Hankosky et al., 2018a, 2018b; Johansen and McFadden, 2017; Kearns et al., 2022; Lin et al., 2024; McFadden et al., 2018; Pena-Bravo et al., 2019; Pittenger et al., 2021; Venniro et al., 2017; Westbrook et al., 2020). Our study achieved the goal of explaining these seemingly inconsistent observations in Table 1.

Interestingly, our data revealed also that saline SA control groups do not represent a homogeneous population. Thus, going forward, caution must be taken when controls are compared with experimental groups in SA experiments.

A major limitation of the study includes the low number of males used in the saline SA group – this led to a cluster that included only females when it should have also included males. This limited our ability to determine if there was a SEX by cluster interaction for the saline SA group. That said, we determined that the MISSING model was not severely challenged by this limitation. Moreover, in our recent report validating the MISSING model for intra-nucleus accumbens core dopamine-related psychostimulant activity (Job, 2024), we also detected a cluster that consisted only of females. We attribute that, and our findings in this study, to sampling errors beyond our control.

We determined that the MISSING model identifies clusters with significant differences that exceeded the differences between biological sex for all variables. The implication is that when we group subjects by biological sex, we may be simultaneously underestimating other important behavioral phenomena. While the mechanisms governing individual DS and sex differences are important in our efforts to understand addictive substances, the current model SABV may limit our ability to understand *both* individual DS and sex differences and may limit progress in both these endeavors. The MISSING model gives us a new approach for SABV which includes comparing males and females within the same clusters so that we can minimize other variables that may confound our understanding of sex differences.

In conclusion, we validate the MISSING model for psychostimulant self-administration paradigm and reveal that it can explain the inconsistencies in the literature regarding the observation of sex differences in METH SA. The MISSING model reveals that observed sex differences in METH SA are not always due to biological sex. Our conclusions represent a paradigm shift from the *status quo* and may significantly advance the field of sex differences research.

## Acknowledgements

The authors wish to acknowledge Dr. Jean Lud Cadet in whose laboratory MOJ conducted the behavioral experiments. Both authors (BHS and MOJ) contributed to data analysis and to the writing of the manuscript. MOJ designed and conducted behavioral experiments and statistical analysis. The authors also acknowledge Dr. Atul Daiwile and Mr. Michael Chojnacki, who contributed to the behavioral experiments. This work was funded by the Department of Health and Human Services/National Institutes of Health/National Institute on Drug Abuse/Intramural Research Program, Baltimore, MD, USA [grant -DA000552]. This work was also supported by the Francis Lax Fund for Faculty Development at Rowan University. This work was also supported by startup funds from Rowan University, Camden, New Jersey.

